# Low-frequency alternating current stimulation rhythmically suppresses stimulus-induced gamma-band oscillations in visual cortex and impairs perceptual performance

**DOI:** 10.1101/230656

**Authors:** Jim D. Herring, Sophie Esterer, Tom R. Marshall, Ole Jensen, Til O. Bergmann

## Abstract

Alpha oscillations (8-12 Hz) are hypothesized to rhythmically gate sensory processing, reflected by activity in the 40-100 Hz gamma band, via the mechanism of pulsed inhibition. We applied transcranial alternating current stimulation (TACS) at individual alpha frequency (IAF) and flanking frequencies (IAF-4 Hz, IAF+4 Hz) to the occipital cortex of healthy human volunteers during concurrent magnetoencephalography (MEG), while participants performed a visual detection task inducing strong gamma-band responses. Occipital (but not frontal) TACS phasically suppressed stimulus-induced gamma oscillations in the visual cortex and impaired target detection, with stronger phase-to-amplitude coupling predicting behavioral impairments. Frontal control TACS ruled out retino-thalamo-cortical entrainment resulting from (subthreshold) retinal stimulation. All TACS frequencies tested were effective, suggesting that visual gamma-band responses can be modulated by a range of low frequency oscillations. We propose that TACS-induced cortical excitability fluctuations mimic the mechanism of pulsed inhibition, which mediates the function of alpha oscillations in gating sensory processing.

## Introduction

Cortical oscillations and their cross-frequency interaction constitute important mechanisms for the organization of neuronal processing. Alpha-band oscillations (8 – 12 Hz) are hypothesized to rhythmically gate information flow in the brain via the pulsed inhibition of sensory processing, reflected by local gamma-band oscillations (40 – 100 Hz) (Klimesch et al., 2007; Jensen and Mazaheri, 2010). Primarily, we aimed to test the specific hypothesis that the well-described stimulus-induced increase in gamma-band power in the visual cortex, associated with bottom-up visual processing (Bastos et al., 2015; Fries, 2015), can be actively modulated by the phase of slower oscillations, particularly in the alpha band. While correlational data from MEG studies in humans (Osipova et al., 2008) and intralaminar recordings in monkeys (Spaak et al., 2012) has revealed coupling between alpha phase and gamma amplitude, the causal role of alpha oscillations in modulating gamma-band power remains unresolved. We therefore applied transcranial alternating current stimulation (TACS) at individual alpha frequency (IAF) to the visual cortex (Oz-Cz montage) in human volunteers performing a visual detection task to mimic the impact of alpha phase-related cortical excitability fluctuations on endogenous gamma activity during visual stimulus processing. A second goal of this study was to test (i) whether TACS is capable of modulating behaviorally relevant neuronal activity in the human brain at commonly used stimulation intensities, an assumption recently called into question by modelling work (Opitz et al., 2016) and cadaver studies (Underwood, 2016), and (ii) whether its effect can be attributed to transcranial as opposed to mere retinal stimulation (Schutter, 2015). While simultaneous TACS-EEG recordings (Helfrich et al., 2014a; Helfrich et al., 2016) are limited by the spatial interference of stimulation and recording electrodes, both affixed to the scalp, the combination of TDCS/TACS and MEG (Soekadar et al., 2013; Neuling et al., 2015; Witkowski et al., 2015; Marshall et al., 2016) allowed us to transcranially impose oscillating currents on the visual cortex, while assessing stimulus-induced gamma power modulation in the visual cortex directly underlying the TACS electrodes (Oz-Cz montage). Using a combination of spatial filtering and TACS artifact suppression techniques, we extracted gamma-band oscillatory signals from the visual cortex during TACS and estimated cross-frequency TACS-phase-to-gamma-amplitude-coupling. To control for the potential impact of electrical stimulation of the retina and resulting retino-thalamo-cortical entrainment, we also applied TACS with a frontal stimulation montage (Fpz-Cz). To further assess the frequency-specificity of TACS-phase-gamma-amplitude-coupling, we applied TACS at two flanker frequencies of IAF -4 Hz and IAF +4 Hz. We hypothesized that occipital TACS (but not frontal control TACS) at alpha frequency should impose rhythmic excitability fluctuations in the visual cortex, mimicking the effects of spontaneous alpha oscillations. Occipital TACS was therefore expected to cause a general decrease and, more specifically, a rhythmic suppression of visual stimulus-induced gamma power, and consequently a reduction of bottom-up visual stimulus processing and associated perceptual detection performance.

## Results

Participants performed a forced-choice visual discrimination task in which they had to report the rotation direction of a foveally presented asterisk inside an inward moving high-contrast grating (Figure 1A), known to produce a pronounced gamma oscillatory response in early visual cortex (Hoogenboom et al., 2006b). During each trial, we applied TACS in either a visual (Oz-Cz) or a frontal (Fpz-Cz) montage and at either of three frequencies (i.e., IAF -4 Hz, IAF, +4 Hz; Figure 1B,C) while recording ongoing brain oscillatory activity using whole-head magnetoencephalography (Figure 1). Sham trials were intermingled as TACS-free reference epochs. Data from 15 of the 17 participants is reported here, as two showed no detectable gamma band response even in TACS-free Sham trials.

**Figure 1.**
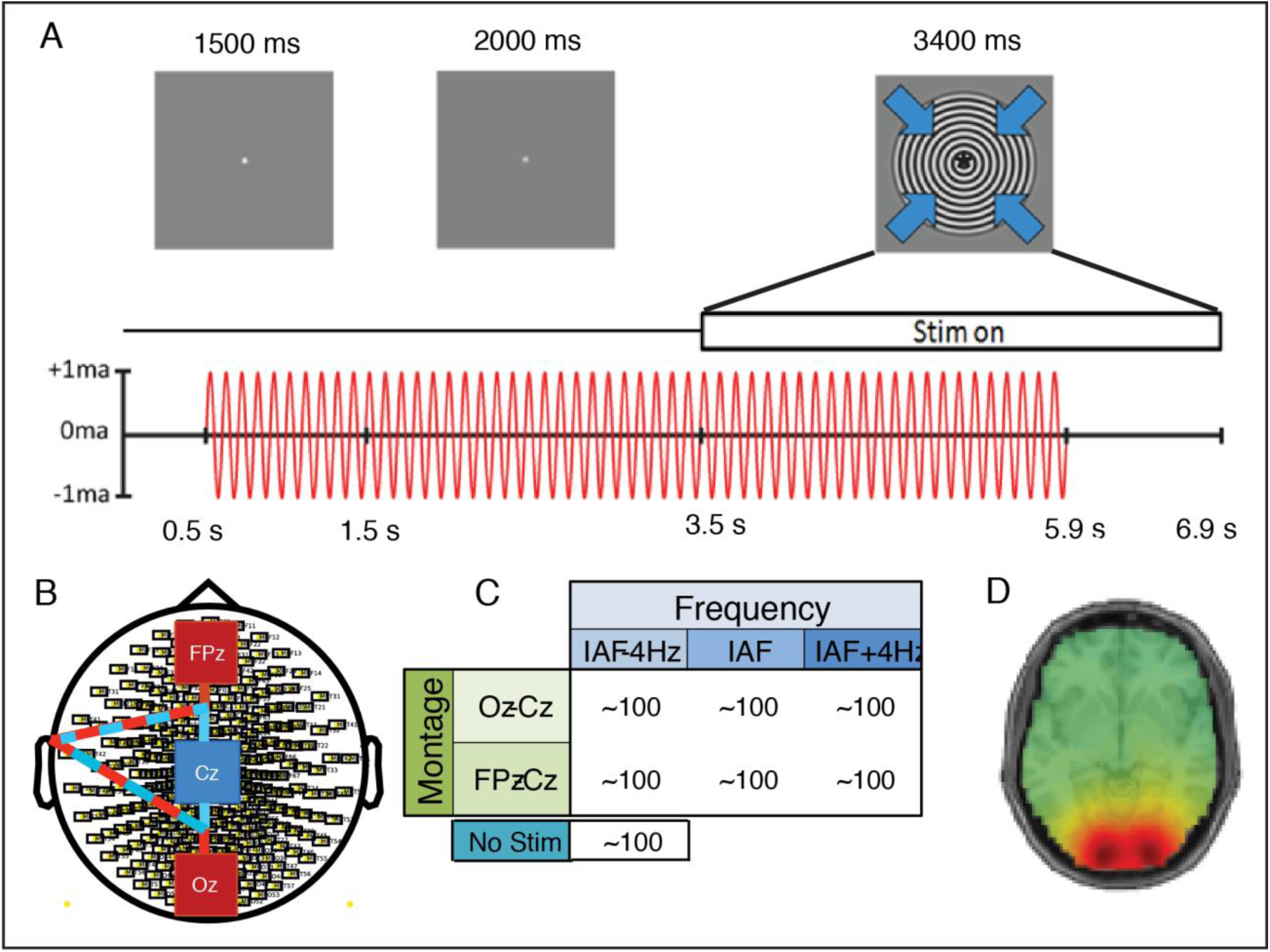
Experimental paradigm and setup. (**A**) Timeline of a single trial. Participants fixated a small white dot in the center of the screen and were allowed to blink, until 1500 ms later the white dot turned grey to indicate the end of the blink period. At 3500 ms an inward-moving grating appeared around the fixation dot, which contained a slowly rotating asterisk in its center. Participants had to report the direction of rotation, by button-press, as soon as the visual stimulus disappeared at 5900 ms and before the next trial started at 6900 ms. TACS was turned on 500 ms into the blink period and turned off 2400 ms after visual stimulus onset and 1000 ms before visual stimulus offset, thus lasting for 5400 ms each trial. (**B**) TACS electrode montage. TACS was applied via three 5 x 5 cm rubber electrodes attached in a dual-montage setup: an occipital montage with electrodes located at Oz and Cz, and a frontal montage with electrodes at Fpz and Cz, with electrode Cz used in both montages. The cables connected to the electrodes were twisted at the shortest possible distance and lead left-ward away from the head towards the shoulder and out of the MEG helmet. (**C)** Experimental design matrix. Seven different trial conditions were pseudorandomly intermingled: 2 montages (frontal, occipital) x 3 frequencies (IAF, IAF -4 Hz, IAF + 4 Hz) plus one stimulation-free condition), with ~100 trials per condition, i.e. ~700 trials in total. Due to limitations in total stimulation duration per day (defined by the local ethics committee) the experiment was split into two sessions of ~350 trials each, separated by at least 1 day. (**D**) Group average topography of stimulus-induced gamma-band power in source space (DICS frequency domain beamforming). Virtual channels were extracted (LCMV time domain beamforming) from the 10 voxels in visual cortex showing the highest relative increase in gamma-band power from baseline.

### Occipital TACS suppressed average gamma power

Gamma power was extracted at individual’s gamma peak frequency (see Methods; see Figure 1D for group average gamma power). Average gamma peak frequency was 57.6 Hz ± 8.5 Hz (mean ± SD). In all conditions, a significant increase in gamma-band power was observed during visual stimulus presentation (one-sample t-tests, all p < 0.001; Table S1). Importantly, visual stimulus-induced gamma responses differed between TACS montages as revealed by the main effect of a *montage* (occipital, frontal, sham) x *frequency* (IAF-4, IAF, IAF+4 Hz) rmANOVA (F_1.3,18.9_ = 26.08; p < 0.001), in that occipital TACS caused a stronger suppression of the gamma response than frontal TACS (paired-samples t-test averaged across TACS frequencies: t14 = -9.15, p_bonf_ < 0.001; see Figure 2A, S2, and S3). In fact, only occipital TACS (t_14_ = -7.59, p_bonf_ < 0.001) but not frontal control TACS (t_14_ = -0.40, p_bonf_ = 1) caused a decrease in gamma power compared to Sham trials, excluding a confound by retinal stimulation. There was no significant main effect for TACS frequency (F_2,28_ = 2.80, p = 0.08) but a significant interaction between montage and frequency (F_4,56_ = 4.38, p = 0.004), driven by occipital TACS causing a larger suppression of gamma-band power for the IAF-4 Hz compared to IAF + 4 Hz conditions (t_14_ = -4.18, p = <0.001), whereas frequencies did not differ for frontal TACS or Sham (all p > 0.1).

**Figure 2.**
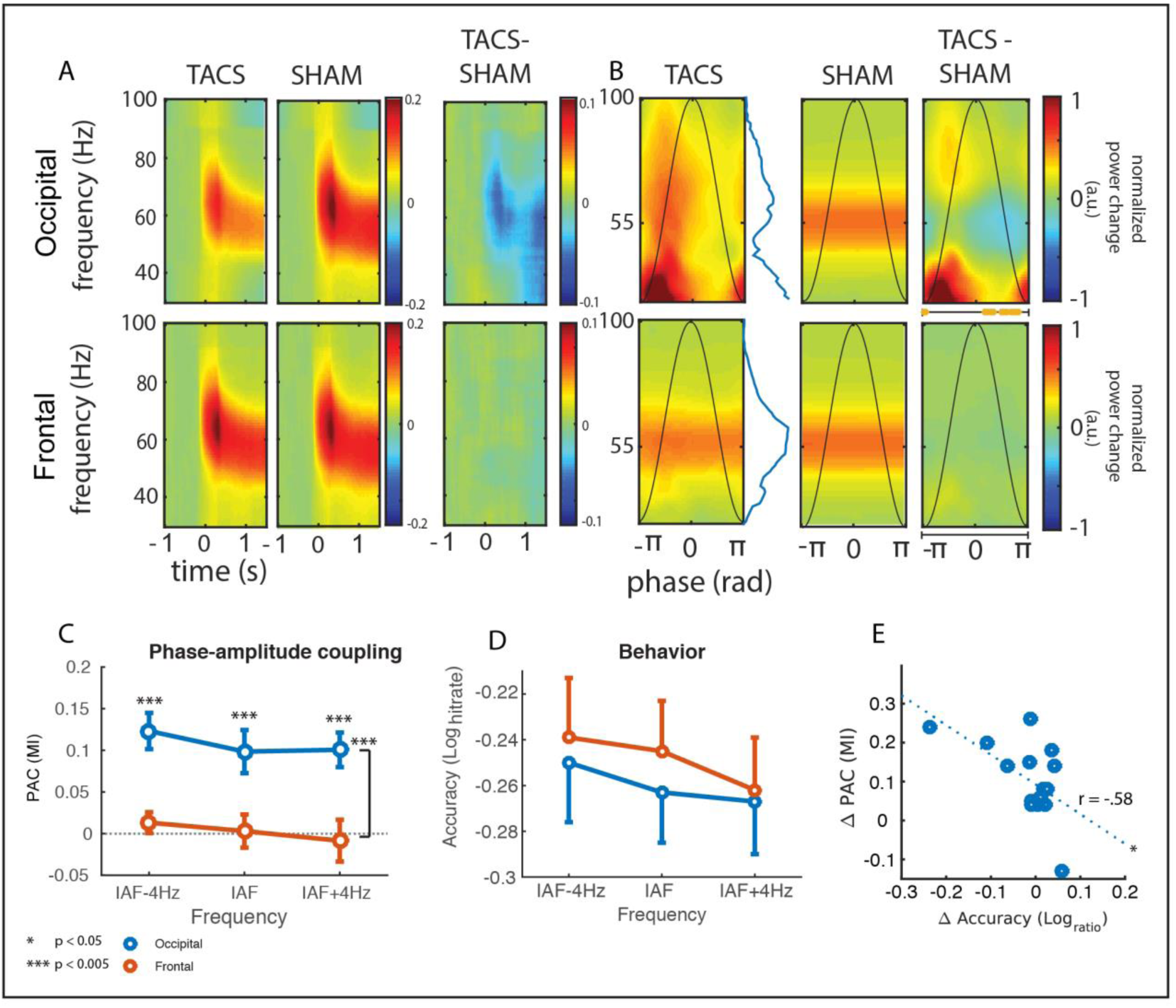
Occipital TACS rhythmically suppressed stimulus-induced gamma oscillations. **(A)** Time-frequency representations (TFR) of oscillatory power timelocked to visual stimulus onset for IAF TACS (TACS) vs. TACS-free trials (Sham) (see Figure S2 for separate analyses of different TACS frequencies). Occipital TACS caused a stronger suppression of the gamma response than frontal TACS (p < 0.001), and only occipital but not frontal TACS decreased gamma power compared to Sham trials (all p < 0.001; Table S2). **(B)** TFRs as in A, but segmented and averaged timelocked to TACS peaks and with the x-axis normalized to phase-angles in radians (see Figure S4 for separate analyses of all frequencies). Inserted curves represent the TACS cycle. Occipital TACS phasically suppressed stimulus-induced gamma-band activity relative to Sham TACS (orange bars underneath TFR indicate significant difference from zero, pFDR < 0.05), whereas frontal TACS did not show any PAC relative to Sham. **(C)** Phase-amplitude coupling (PAC, as indexed by the ‘modulation index’, MI; Tort et al., 2006) between the phase of TACS and the amplitude of the stimulus-induced gamma power during visual stimulus presentation (after subtraction of PAC during TACS in pre-visual-stimulus baseline and surrogate PAC to control for the potential impact of residual TACS artefacts on PAC). PAC was larger for occipital TACS (blue) than for frontal TACS (orange) and for surrogate data for all TACS frequencies (all p < 0.005), whereas frontal TACS did not differ from surrogates (p > 0.3). See Figure S4 for periods before and during visual stimulation. **(D)** Accuracy on the rotation-detection task was reduced for occipital compared to frontal TACS (p < 0.05), irrespective of TACS frequency. **(E)** The more the individual TACS-phase-gamma-amplitude-coupling differed between occipital and frontal TACS, the stronger was the individual performance decrease in rotation detection for occipital relative to frontal TACS (r16 = -0.58; p < 0.05).

### Occipital TACS phase rhythmically modulated gamma power

To test whether the net suppression actually resulted from a phasic modulation, or more specifically, a rhythmic suppression of gamma-band power by TACS phase, we calculated TACS peak-locked TFRs (Figure 2B; Figure S3). Visually induced gamma-band power was observed to decrease at particular points in the phase cycle, but not to increase at any point (p_fdr_ < 0.05). In addition, there was a rhythmic increase around 40 Hz, as well as between 70 – 100 Hz for occipital, but not for frontal TACS between – pi and 0 (p_fdr_ < 0.05). Although activity in the 40 Hz range is also visible in the non-peak-locked TFRs (Figure 2A), it is outside the range of the main gamma-band response. To more formally quantify the strength of phasic gamma power modulation, we assessed phase-amplitude coupling (PAC) between TACS phase and gamma-band power in the visual cortex by calculating Tort’s Modulation Index (MI, Tort et al., 2010) both before and during visual stimulus presentation. We calculated the MI for each TACS condition at the individual peak-gamma frequency. For comparison, montage- and frequency-specific surrogate samples were created by phase-shifting the TACS signal from the respective TACS condition. PAC was significantly larger for occipital than for frontal TACS, as reflected by a significant main effect of montage (Figure 2C) during visual stimulation (F_1,14_ = 64.53, p < 0.001), but neither a main effect of frequency (p > 0.2) nor an interaction (p > 0.6). Importantly, this effect remained significant (F_1,14_ = 17.06, p < 0.001) after correcting for potentially spurious phase-amplitude coupling. To this end we subtracted PAC values calculated on the baseline period, which included TACS but lacked the visual stimulus-induced gamma response (peri_TACS_-pre_TACS_). In addition, we calculated the same difference in PAC values for the surrogate data (perisurrogate-presurrogate), the result of which was subtracted from the respective TACS difference ((peri_TACS_-pre_TACS_)-(peri_surrogate_-pre_surrogate_)). In fact, only occipital (t_14_ = 5.23, p < 0.001) but not frontal TACS (t_14_ = 0.168, p > 0.8) showed significant PAC during visual stimulus presentation relative to baseline and relative to the respective effect in the PAC surrogates. Thus, ruling out retinal entrainment and spurious PAC due to residual artifacts, occipital TACS did indeed produce a phasic modulation of stimulus-induced gamma power that was comparable in strength across stimulation frequencies.

### Occipital TACS decreases rotation discrimination

Participants correctly identified the rotation direction in 78% of trials (SEM = 2%) with an average reaction time of 409 ms (SEM = 11 ms), for correct trials. There was a small, but significant decrease in hit rate for occipital (77.98% ± 2.66%) compared to frontal TACS (78.33% ± 2.26%) when taking into account the participant’s baseline performance in Sham trials as a covariate in a 2 x 3 repeated-measures ANCOVA (montage x TACS frequency; see Figure 2D). We observed a main effect of montage (F_1,15_ = 5.81, p < 0.05) and an interaction between montage and baseline performance in the Sham trials (F_1,15_ =11.69, p < 0.01). The interaction indicates that subjects performing better in the rotation discrimination task during Sham also showed stronger TACS-related impairment than weakly performing subjects, a relationship that is also reflected by the correlation between performance during Sham trials and performance reduction during occipital TACS relative to frontal control TACS (r_16_ = 0.66, p < 0.01). Importantly, the effect of TACS montage on behavioral performance was correlated with the effect of TACS montage on PAC (r_13_=-.5753, p < 0.05; Figure 2E). In other words, subjects in which occipital TACS caused a stronger rhythmic suppression of stimulus-induced gamma-band activity (compared to frontal control TACS) also suffered from stronger performance impairment in the rotation discrimination task for occipital TACS (compared to frontal control TACS) (Figure 2E).

## Discussion

The aim of this study was to test a core prediction of the *pulsed inhibition hypothesis* (Klimesch et al., 2007; Jensen and Mazaheri, 2010), namely that the phase of slower oscillations, particularly in the alpha band, modulates the bottom-up processing of sensory information, which is reflected by stimulus-induced gamma-band oscillations (Bastos et al., 2015; Fries, 2015), and thereby affects perceptual performance. In line with this notion, we found that TACS to the occipital cortex (but not frontal control TACS) rhythmically suppressed visual stimulus-induced gamma-band power and that the degree of this suppression predicted the reduction in visual detection performance. The rhythmic fluctuations in visual cortex excitability imposed by TACS therefore seems to (by means of entrainment or otherwise) mimic the mechanism of *pulsed inhibition* proposedly constituted by spontaneous alpha oscillations, thus recreating the functional effects of alpha oscillations on sensory processing.

### TACS rhythmically suppresses visual-induced gamma power

The net suppression of visually induced gamma power by occipital TACS (Figure 2A) ties in well with the observed phase-specific decrease (but not increase) in gamma power (Figure 2B), suggesting that the corresponding excitability fluctuations imposed on the visual cortex by occipital TASC may be asymmetric, just as has been proposed for spontaneous alpha-band oscillations (Jensen and Mazaheri, 2010; Schalk, 2015). Notably, TACS at neighboring frequencies 4 Hz slower or faster than individual alpha frequency also caused a net suppression (Figure 2A) and rhythmic modulation (Figure 2C) of stimulus-induced gamma responses (with the lowest TACS frequency even causing the strongest net suppression). There are three potential explanations for the lack of frequency-specificity with respect to phase-amplitude coupling: Firstly, TACS at neighboring frequencies 4 Hz away from individual peak frequency may have still been able to entrain ongoing spontaneous alpha oscillations (Helfrich et al., 2014a; Antal and Herrmann, 2016). This would be in line with the targeted oscillatory network showing resonance properties reflecting an Arnold Tongue, according to which stimulation slightly off the endogenous frequency can still cause entrainment if applied at higher intensities (Ali et al., 2013). Secondly, TACS at neighboring frequencies may have entrained high theta band and low beta band oscillations instead, both of which phasically modulate gamma power in the visual cortex (Bastos et al., 2015; Fries, 2015). Thirdly, transcranial currents may have directly imposed excitability fluctuations at stimulation frequency on relevant visual cortex neurons by merely mimicking the functional aspect of endogenous low-frequency neuronal oscillations without the ‘entrainment’ of an already ongoing endogenously generated oscillation. While the current study cannot rule out any of these explanations, the latter explanation may be favorable as it requires the least assumptions. Our findings are complemented by work from Helfrich et al. (2016), who reanalyzed an earlier TACS-EEG dataset (Helfrich et al., 2014b) and found increased cross-frequency coupling between spontaneous alpha oscillations (8-12 Hz band) and task-related gamma band activity during 10 Hz TACS compared to sham. While these results support the idea that TACS at 10 Hz can facilitate alpha-band to gamma coupling, they did not resolve at that time whether gamma was entrained at the precise TACS frequency, and whether retinal entrainment may have been involved, two issues we explicitly addressed in this study.

### TACS related gamma modulation is behaviorally relevant

Occipital TACS compared to frontal control TACS caused a small but significant impairment in visual detection performance, which can, importantly, not be attributed to retinal stimulation. Beyond that, subjects showing a stronger TACS related modulation of gamma power also showed stronger drops in visual accuracy. Thus, as predicted by the *pulsed inhibition hypothesis*, phasic suppression of stimulus-related gamma oscillations suppressed bottom-up neuronal processing and thus impaired perception. Importantly, the behavioral relevance of TACS-related neuronal activity modulations already shows that the observed neuronal effects cannot be attributed to residual artifacts.

### TACS effects cannot be explained by subthreshold retinal entrainment or residual artifacts

One of the key challenges in studies using TACS is not only the current flow to unintended brain regions, but also to extracranial neuronal structures, such as the retina, which is particularly sensitive to stimulation; TACS easily excites the retina and induces retinal phosphenes, i.e., a sensation of flickering light at stimulation frequency (Schutter, 2015). Importantly, even stimulation intensities below phosphene threshold may entrain visual cortex activity via the retino-thalamo-cortical pathway, explaining why even subconscious intermittent photic stimulation can produce cognitive effects (for a review see Schutter, 2015). To exclude this potential confound, we included a control montage (Fpz-Cz) with the frontal electrode even closer the eyes and matched the stimulation intensity on the retina for both montages by independently adjusting it to 80 % of the subjects’ individual phosphene threshold. This ensured that (i) no retinal phosphenes were induced throughout the experiment, and (ii) the amount of effective current reaching the retina was comparable between montages, while only the occipital montage exerted a direct transcranial impact on the visual cortex. Indeed, we did not observe any effect of the frontal control TACS on either behavior, or on phase-amplitude coupling with gamma band power, or on the relationship between both. This demonstrates that the behaviorally relevant modulatory effects of occipital TACS observed in the current experiment are not explained by indirect retino-thalamo-cortical but rather by direct transcranial cortical entrainment.

A second challenge for TACS studies in combination with EEG or MEG are the enormous stimulation artifacts that contaminate the recordings, particularly at the stimulation frequency but also at its harmonics, as well as at heartbeat- and respiration-related side bands (Noury et al., 2016; Noury and Siegel, 2017). While beamformer spatial filter techniques (like the one used in the current study) can considerably attenuate the artifact when correctly parameterized (Neuling et al., 2017), they may not be able to remove it completely due to its non-linearity (Noury et al., 2016) and mathematical constraints related to (un-)correlated sources (Mäkelä et al., 2017). In the current study, we deliberately circumvent most of these issues by focusing entirely on the effects of low-frequency TACS on high-frequency stimulus-induced gamma power. Importantly, we applied identical artifact removal procedures to all TACS montages, TACS frequencies, and created respective frequency-specific Sham conditions (see Methods for details). Moreover, TACS periods before and during visual stimulation (containing the same amount of residual TACS artifacts) were contrasted to determine induced gamma response (see Methods for details), an approach for which exists consensus that it effectively controls for residual TACS artifacts (Neuling et al., 2017; Noury and Siegel, 2017). We are thus confident that none of the reported findings can be attributed to TACS artifacts systematically confounding power or PAC estimates.

### General implications for TACS-MEG/EEG research

Given the increasing use of TACS and TDCS for the non-invasive modulation of neuronal activity and cognitive function on the one hand, and the recent criticism regarding replicability (Horvath et al., 2014, 2015), effectiveness of commonly applied current intensities (but see Opitz et al., 2016; Underwood, 2016; Opitz et al., 2017), and confounds by peripheral (e.g., retinal) entrainment (Schutter, 2015) on the other hand, our findings have important implications reaching beyond the specific research question of the present study. Numerous studies (e.g., Pogosyan et al., 2009; Joundi et al., 2012; Neuling et al., 2012; Brittain et al., 2013; Helfrich et al., 2014a; Cecere et al., 2015; Alekseichuk et al., 2016) have provided compelling behavioral evidence that TACS does affect human brain function. However, due to the massive presence of TACS artifacts in EEG/MEG recordings (see above), direct assessment of a TACS-related phase-dependent modulation of neuronal activity at the stimulation frequency is inherently problematic (Bergmann et al., 2016). In contrast, by investigating TACS-phase-to-gamma-power-coupling, we could demonstrate that TACS is indeed able to rhythmically suppress visual stimulus-induced gamma oscillations in a behaviorally relevant manner, while ruling out retinal entrainment or residual artifacts as alternative explanation. This implicates that, if adequately controlled, TACS-MEG at common stimulation intensities is a powerful tool to study and manipulate oscillatory brain activity and behavioral performance non-invasively in humans.

## Methods

### Participants

All participants were recruited from a database of the Radboud University Nijmegen. In total, 17 participants (5 males, 12 females, age: 24.3 ± 0.7 (mean ± SEM)) with normal, or corrected-to-normal vision by contact lenses only, were included in the study. All participants conformed to standard inclusion criteria for MRI, MEG, and TACS. Written informed consent was obtained prior to start of the experiment according to the Declaration of Helsinki. The study was approved by the local ethics committee. Participants were financially compensated at 10 Euros per hour. Data are reported from 15 participants; two participants had to be excluded since no visual stimulus-induced gamma-band response could be detected even during Sham periods.

### Procedure

To gather a sufficient amount of trials, participants took part in two experimental sessions on two separate days, but with all experimental conditions tested in each session. A structural MRI was obtained on a separate day. In the beginning of the first session, participants were familiarized with the experimental task. Otherwise, both experimental sessions followed the same procedures: After TACS and ECG electrodes were applied, participants were familiarized with the stimulation, and individual stimulation intensity was determined. Then, participants were seated in the MEG, and four minutes of resting state data were collected (two minutes eyes-open, two minutes eyes-closed) before they performed a rotation-detection task in blocks of 10 minutes with short breaks in between. Total MEG time was 90 min per session.

### Rotation detection task

Participants performed a rotation detection task in which they had to indicate by button-press the rotation direction of an asterisk in the center of an inward moving high-contrast grating (Figure 1A). Participants were instructed to fixate a white dot on grey background in the center of the screen. Each trial started with a 1.5 s period in which participants were allowed to blink. Participants were asked to refrain from blinking as soon as the fixation dot turned grey. A ‘baseline’ period of two seconds followed after which an inward-moving (0.8 degree/second) black-and-white high contrast grating with concentric circles (2.5 cycles/degree) appeared on screen covering 8 degrees of visual angle (adapted from Hoogenboom et al., 2006a) for 3.4 seconds. In the center of the inward moving grating an asterisk was present that slowly rotated either clock-wise or counter-clock wise. The rotation rate was continuously updated after each trial using an adaptive-staircase procedure (Watson and Pelli, 1983) so that participants were roughly 80% correct in detecting rotation direction. The goal of the task was to keep the participants fixated and assure a stable level of attention throughout the experiment, as well as to assess TACS effects on foveal detection accuracy.

### TACS

TACS was applied using a battery-driven NeuroConn DC+ stimulator (neuroConn GmbH, Ilmenau, Germany) connected to three 5 x 5 cm conductive, non-ferromagnetic rubber electrodes that were attached to the scalp following the international 10-20 system, creating an occipital montage (Oz-Cz) and a frontal montage (Fpz-Cz), which shared electrode Cz (Figure 2B). The surface area of the scalp was thoroughly cleaned using alcohol and Nuprep skin preparation gel (Weaver and Company, Aurora, CO, USA). Then electrodes were attached using conductive Ten20 paste (Weaver and Company, Aurora, CO, USA) ensuring no paste was applied outside of the contact area of the electrodes (Marshall et al., 2016). Impedances were kept below 5 kOhm. Due to the stickiness of the paste no further mounting aids were required. Electrode cables were connected in a manner that minimized the size of the current loop formed on the scalp. Cables for the Oz-Cz montage were connected to the anterior side of electrode Oz, and to the posterior side of electrode Cz. Cables for the Fpz-Cz montage were connected to the posterior side of electrode FPz, and the anterior side of electrode Cz. Switching between montages was handled by galvanically isolated switchbox that was controlled by the stimulation PC.

Several steps were taken to minimize the artifacts produced by the presence of additional material in the magnetically shielded room (MSR). First, all electrodes and cables, including connectors, were checked for ferromagnetic properties by moving the items inside the helmet while inspecting the effects on the MEG signal (several rubber electrodes had to be dismissed as they produced artifacts in the MEG signal). Second, a CAT6 electronically shielded cable connected the electrodes to the stimulator, which was located outside the MSR. The cable was fixed to the chair in the MSR to minimize movement. Third, the cables attached to the scalp were twisted for each montage to keep the resulting current loop as small as possible. The cables attached to the scalp ran left of the subject, fixed to the shoulder, downwards towards the chair, and away from the helmet.

Stimulation frequency was adjusted per participant based on the individual alpha frequency. To this end, the individual alpha frequency (IAF) was determined, with a resolution of 0.2 Hz, at the beginning of each session as the peak frequency of the difference in power spectra between an eyes-closed and eyes-opened resting state session (10.31 ± 0.41 Hz, mean ± SD across subjects). During the experiment, stimulation frequency was set for each trial to either IAF -4Hz, IAF, or IAF +4 Hz (Figure 2C). These two flanker frequencies at 4 Hz below and above IAF were chosen to explore the frequency specificity of the stimulation while still being sufficiently close to the alpha-band not to target neighboring functionally relevant frequency bands (i.e., the theta- or beta-band).

Stimulation intensity was titrated at the beginning of each session to 90% of the individual retinal phosphene threshold, separately per montage (peak-to-peak TACS amplitude for Oz-Cz: 963 ± 319 μA and Fpz-Cz: 231 ± 114 μA; mean ± SD, range for Oz-Cz: 450 – 1750 μA, for FPz-Cz: 50 – 550 μA). Retinal phosphene threshold was determined by increasing the current strength of TACS at IAF from 100 mA in steps of 100 mA until the participants started to perceive retinal phosphenes. Note that stimulation intensity was adjusted per montage, since the goal of the frontal montage was to control for retinal stimulation effects by increasing the proximity to the retina, while decreasing the proximity to the occipital cortex, ensuring that the current effectively stimulating the retina was comparable between montages. The frontal montage was not meant to effectively stimulate the frontal cortex (in which case higher stimulation intensities would have been required). The choice of subthreshold intensity for both montages ensured that no visual phosphene perception interfered with (i) the transcranial stimulation effects, (ii) the visual stimulus-induced gamma responses, and (iii) the detection task performance.

During each trial (except for Sham trials), TACS was applied for ~5.4 s at one of the three frequencies (IAF -4Hz, IAF, or IAF +4 Hz) and via one of the two montages (occipital Oz-Cz or frontal Fpz-Cz) (Figure 1C). Stimulation started 1 s before the baseline period to allow for a build-up of potential entrainment effects and continued throughout the 2 s baseline period into the visual stimulus presentation period. The stimulation was turned off 2.4 seconds into the visual stimulation period after completing a full number of cycles at the particular stimulation frequency. In total, 700 trials (600 TACS trials + 100 Sham trials) were acquired per subject, distributed over two sessions (resulting in a total stimulation time of 27 min per session.), and pseudo-randomized in order (i.e., all seven experimental conditions randomly applied within each consecutive block of seven trials) to ensure that an equal amount of trials of each condition was completed after every session and that there would be no more than two direct repetitions of the same TACS condition.

### MEG data acquisition

Whole-head MEG was recorded using a 275-channel axial gradiometer CTF system (CTF MEG systems, VSM MedTech Ltd.) sampling at 1200 Hz, with a hardware low pass filter at 300 Hz. Head localization coils were placed on the nasion, and in the left- and right-ear canals. The position of the head was recorded at the beginning of the experiment and was monitored, and adjusted during breaks using online head-position tracking if head motion exceeded 3 mm (Stolk et al., 2013). Eye-tracking was conducted throughout the experiment using an EyeLink 1000 eyetracker (SR Research Ltd, Ottawa, Canada), sampling at 2 kHz. Electrocardiogram (ECG) was recorded using two electrodes in a bipolar montage placed on the left collarbone, and right hip.

### Data Analysis

#### Preprocessing

Data analyses was conducted using the FieldTrip toolbox (Oostenveld et al., 2011) and custom Matlab scripts for Matlab 2014b (Mathworks, Nattick, USA). First, trials that included blinks during the baseline-or visual stimulation period were detected by bandpass filtering the horizontal, and vertical motion eye-tracker channels between 1 and 15 Hz (4^th^ order, two-pass, Butterworth). Trials that exceeded a z-score of 5 were rejected. Second, trials that included SQUID-jumps were detected by first high-pass filtering the data at 30 Hz (4^th^ order, two-pass, Butterworth) to attenuate the stimulation artifact. Trials of which the first-order temporal derivative exceeded a z-score of 25 were rejected. This resulted in an average of 9% ± 8% (Mean ± SD) rejected trials per subject. The data were down-sampled to 600 Hz after epoching into trials from -3.4 to 3.6 seconds after onset of the gamma-inducing visual stimulus.

### DICS beamforming

A single-shell head model (Nolte, 2003) was created from the individual MRIs. Next, an equally-spaced grid with 0.5 mm^3^ based on a standard MNI template MRI with 0.1 mm^3^ resolution was created. This template grid was warped to each subject’s individual anatomy to easily average and compare voxels across subject’s. Then, a spatial filter was designed to maximize the sensitivity to the expected gamma-band response produced by the visual stimulation. To this end, we selected the trials without stimulation and epoched them into a baseline period of -2.0 to -0.001 s and an activation period from 0.4 to 2.399 s after visual stimulus onset, thus creating two epochs of exactly 2.0 s length. After removing linear trends, we calculated the cross-spectral density (CSD) matrix for both baseline and activation epochs, as well as for both epochs combined, at 60 Hz with 15 Hz frequency smoothing using a multi-taper approach, thus resulting in an analyzed frequency band of 45 – 75 Hz. A common spatial filter was calculated using a Dynamic Imaging of Coherent Sources (DICS) beamformer (Gross et al., 2001) on the CSD of the combined baseline and activation data using 5 % regularization. Note that the spatial filter was calculated on TACS-free Sham trials. The resulting spatial filter was then applied to the activation and baseline data separately. To find the voxels showing the maximal increase in gamma-band power in response to visual stimulation, the relative gamma-power change from baseline was calculated by dividing for each voxel in source space the absolute change from baseline by the baseline gamma power.

### LCMV virtual channels

To enhance sensitivity to the visually induced gamma-band response, we used Linear Constrained Minimum Variance (LCMV) beamforming (Van Veen et al., 1997) to extract virtual channel time courses from those voxels that showed the strongest gamma-band power increase from baseline in each participant (Figure 1D) and were located inside the visual cortex mask of the AAL atlas (Tzourio-Mazoyer et al., 2002), including all striate and extra-striate regions (method adapted from Marshall et al., 2015). From these data, we created a new grid with 10 voxels. After bandpass-filtering (40 – 70 Hz) to maximize spatial filter sensitivity to the gamma-band, the covariance matrix was calculated for epochs from -2.3 to 2.3 s relative to visual stimulus onset on Sham trials, and spatial filters were calculated using 5% regularization. The raw sensor-level data was multiplied by the resulting spatial filters to obtain virtual channel time courses for each of the 10 voxels in the grid. For each subsequent analysis, we first analyzed each of the 10 time courses separately before averaging the results per subject.

### FFT interpolation

The main focus of the study was the effect of TACS in the alpha frequency-range on visually-induced gamma-band oscillations. Therefore, our original strategy was to ignore the artifact-loaded signal at the stimulation frequency itself and only analyze the gamma-band power modulation with respect to the known TACS phase. However, while TACS was applied at lower frequencies (range: 5 – 16 Hz) and thus well outside the stimulus-induced gamma-band of interest (i.e., 45 –75 Hz), the magnitude of the TACS artifact in the MEG signal was orders of magnitude larger than the magnetic fields produced by the brain, and the higher harmonics of the TACS frequency still affected the gamma-band frequencies of interest (Supplementary Figure S1C). TACS artifacts and their harmonics could not be sufficiently suppressed by bandstop filters alone. LCMV spatial filters have previously been used to extract the brain signal of interest while attenuating the TCS artifact due to the suppression of correlated sources (e.g. Neuling et al., 2015; Marshall et al., 2016; Neuling et al., 2017), although a full suppression is mathematically impossible (Mäkelä et al., 2017). Unfortunately, we could not follow this approach, since LCMV spatial filter calculation based on TACS trials did not only suppress TACS artifacts, but also the stimulus-induced gamma-band response of interest: In fact, with that procedure, only 3 out of 17 participants still showed a clear gamma-band response during Sham trials, although the gamma-response is known to be very reliable (Hoogenboom et al., 2006a; Scheeringa et al., 2009; Scheeringa et al., 2011). In contrast, with our approach, only 2 out of 17 subjects did not show a visible gamma-band response. We therefore calculated LCMV spatial filters on Sham trials only, with the priority of preserving gamma-band responses in the visual cortex, while still attenuating TACS artifacts, though to a lesser degree. In addition, we employed an FFT interpolation approach (Figure S1C,D) previously used to attenuate line-noise in ECG recordings (Mewett et al., 2001) to effectively suppress the TACS frequency and its harmonics in all conditions. Data were transformed to the frequency domain using a Fast Fourier transform (FFT) with 0.2 Hz frequency resolution (after zero-padding each trial to 5 s). Then, for each TACS trial the magnitude spectrum was interpolated from -1 to 1 Hz around the TACS frequency, and each of its harmonics up until the Nyquist frequency, while the phase spectrum remained intact. The interpolated frequency domain data was then transformed back into the time domain using an inverse FFT. As the effect of overall magnitude attenuation of the signal depended on the TACS frequency, we generated appropriate control conditions by applying the same FFT interpolation approach to copies of the Sham trials. The resulting frequency-specific Sham control conditions thus ensured fair comparisons even if harmonics inside the gamma frequency-band were interpolated.

### Time-Frequency Analysis

To assess visual stimulus-induced gamma-band responses we calculated time-frequency representations (TFRs) of power by means of a sliding window FFT. For each trial, a sliding time window of 500 ms was moved in steps of 20 ms over the entire trial. The sliding time window was multiplied with a sequence of tapers (discrete prolate slepian sequence; dpss) to achieve a frequency smoothing of 10 Hz. Frequencies between 30 Hz and 100 Hz were analyzed in steps of 1 Hz. The data was zero-padded up to 10 seconds to achieve an artificial frequency resolution of 0.1 Hz. The mean and any linear trend were removed prior to calculating the FFT. The gamma-band response is usually best represented as relative change from baseline to account for its comparably low amplitude. During TACS trials, however, any residual noise in the baseline may thereby result in spuriously low ratios. We thus first subtracted the baseline (-1 to -0.2 s) from the activation period, effectively removing any residual frequency- and montage-specific TACS-related artifacts, as well as their heartbeat- and respiration-related modulation (Noury et al., 2016; but see Neuling et al., 2017), which per design were identical in baseline and visual activation periods of the same condition. We then calculated the log-ratio with respect to a common baseline period derived from the average of all Sham trials and thus unaffected by TACS-artifacts (ensuring a fair comparison of the gamma-band response between conditions and relative to Sham trials). An unbiased estimate of individual gamma peak frequency was derived by first calculating, for all conditions, the relative change in gamma power from Sham baseline, fitting a 23^rd^ order polynomial to the average over all conditions, and identifying the largest peak within a range from 40 – 90 Hz using Matlab’s ‘findpeaks’ function. Relative gamma power was extracted per experimental condition from individual peak frequency bins and the time period from 0.5 to 1.5 s post visual stimulus onset and used for all subsequent analyses. Note that the attenuation of gamma power differs between frequency conditions based on the number of TACS harmonics in the gamma-band range that had to be interpolated in the frequency domain for artifact removal. However, the same procedure was applied to respective Sham trials, to guarantee a fair comparison. We used one-sample t-tests to test for significant gamma power responses relative to baseline, and a montage (occipital, frontal, sham) x TACS frequency (IAF-4, IAF, IAF+4 Hz) repeated-measures ANOVAs (with Greenhouse-Geisser correction for non-sphericity where necessary) on the baseline- and surrogate-corrected data, followed by paired-sample t-tests (with Bonferroni correction for multiple comparisons) were appropriate, to compare values between TACS montage and frequency conditions.

### Alpha Peak-Locked TFRs

To assess whether TACS phasically modulated the power of the visually-induced gamma-band responses, we evaluated the gamma-band power dynamics in TACS peak-locked TFRs. To this end, we first calculated TFRs of each trial as described in the previous paragraph, but decreased the size of the sliding time window to 0.1 s to be more sensitive to transient changes in the gamma-band across the TACS cycle. Also, data were normalized per trial to allow robust single-trial assessment of phasic gamma power modulation for subsequent TACS-phase-gamma-amplitude-coupling analyses. To take variations in signal-to-noise ratio into account (e.g., due to residual artifacts), TFRs were z-normalized trial-by-trial by subtracting the mean and dividing by the standard deviation of the trial’s baseline, resulting in an estimate of time-locked power that is relatively robust against noisy trials and extreme values (see Grandchamp and Delorme (2011)).

Next, we detected the peaks of the TACS cycle in the output copy of the TACS signal as provided by the stimulation device, using Matlab’s *findpeaks* function. Peaks were defined as the local maxima on the z-transformed stimulation signal with a minimum width of a quarter cycle of the stimulation signal, a minimum height of 1, and a minimum distance of 0.9 cycles. The data were epoched into segments around each peak with a duration of 4 cycles of the respective TACS frequency. For each TACS frequency, a comparable segmentation was also applied to the Sham trials, using randomly chosen stimulation signals from the respective TACS trials. This resulted in a specific Sham control condition for each TACS frequency, effectively providing a surrogate distribution of gamma power values relative to the TACS cycle. Finally, averages were created for each of the TACS frequencies and respective Sham surrogates. As the number of epochs depends on the number of cycles (being higher for higher frequencies), we applied a random subsampling approach to create unbiased averages. We first determined the stimulation frequency with the smallest number of epochs and then averaged 500 randomly drawn subsamples of that size per condition. To allow direct comparison between TACS frequencies and averaging across subjects (with individualized IAF), we transformed the time-axis to radians by adjusting the step size during TFR calculation accordingly. Importantly, TACS peak-locked TFRs were calculated for both baseline and visual stimulation period. Since the baseline period does not contain any visually-induced gamma-band responses, but may contain residual TACS artifacts, it serves as an excellent control against TACS artifact-related spurious phase-amplitude coupling. Respective Sham TFRs were then subtracted from the TACS peak-locked TFRs, and individual gamma power values were compared for each phase angle (radians) with two-sided one-sampled t-tests against zero using FDR correction for multiple comparisons (Figure 2B and Figure S3).

### Phase-Amplitude Coupling

To more formally quantify and compare the extent to which TACS phasically modulates the gamma-band response, we estimated phase-amplitude coupling (PAC) using Tort’s Modulation Index (MI) by calculating the normalized Kullback-Leibler (KL) divergence of the histogram of TACS phase-binned gamma amplitude to a uniform distribution (Tort et al., 2010). In case of significant PAC, the histogram diverges from a uniform distribution. To this end, the gamma-band amplitude was determined by convolving the virtual channel data between 0.5 and 1.5 s after visual stimulus onset with a 5-cycle moving time window multiplied with a Hanning taper for frequencies from 30 Hz to 100 Hz in steps of 1 Hz, while a 1 second time window was used for estimating the phase of the TACS signal similarly to the gamma-band magnitude (Jiang et al., 2015). The phase-difference between the gamma-band power envelope and TACS signal was subsequently calculated and compared to a uniform distribution using Tort’s Modulation Index (MI). As for TACS peak-locked TFRs, we randomly subsampled the data for each condition 500 times using a sample size equal to the lowest number of trials across conditions, to prevent any bias due to unequal trial numbers between conditions. MIs were calculated for each random subsample and then averaged. As a control, PAC was also estimated for surrogate data, for which the phase-providing TACS signal was randomly phase-shifted to create frequency-specific surrogate PAC values for each TACS condition. We used one-sample t-tests to test for significant PAC after subtracting PAC values at baseline before visual stimulus onset and respective visual-stimulation induced changes in surrogate PAC values. We used repeated-measures ANOVAs (no correction for non-sphericity was necessary) on the baseline- and TACS-corrected data, followed by paired-sample t-tests were appropriate, to compare PAC between TACS montage and frequency conditions.

## Acknowledgements

This work was supported by The Netherlands Organization for Scientific Research, VICI Grant 453-09-002, ALW Open Competition Grant 822-02-011.

## Supplemental Tables and Figures

**Table S1.** *Visual stimulus-induced gamma power*. Table contains mean (± SEM) for visual stimulus-induced gamma power, calculated per condition by subtracting individual gamma power values of the respective ‘baseline’ period before visual stimulation from the ‘activation’ period during of visual stimulation and subsequently dividing those differences by the individual baseline of the Sham condition to normalize for power differences across subjects. Right columns contain t-statistics and p-values for one-sample t-tests against zero. All means were significantly different from zero, indicating an increase in gamma-band power being evident in all conditions relative to a common pre-visual-stimulus baseline as derived from Sham trials.

**One Sample T-Tests**
|  | Mean | SE | t | df | p |
| --- | --- | --- | --- | --- | --- |
| Occipital: IAF - 4 Hz | 0.095 | 0.016 | 5.964 | 14 | < .001 |
| Occipital: IAF | 0.108 | 0.020 | 5.356 | 14 | < .001 |
| Occipital: IAF + 4 Hz | 0.122 | 0.018 | 6.920 | 14 | < .001 |
| Frontal: IAF - 4 Hz | 0.158 | 0.019 | 7.453 | 14 | < .001 |
| Frontal: IAF | 0.152 | 0.020 | 7.490 | 14 | < .001 |
| Frontal: IAF + 4 Hz | 0.156 | 0.020 | 7.823 | 14 | < .001 |
| Sham: IAF - 4 Hz | 0.155 | 0.021 | 8.140 | 14 | < .001 |
| Sham: IAF | 0.157 | 0.020 | 7.448 | 14 | < .001 |
| Sham: IAF + 4 Hz | 0.158 | 0.021 | 7.845 | 14 | < .001 |

*Note.* Student's t-test.

**Table S2.** *Difference of visual stimulus-induced gamma power relative to Sham trials*. Table contains comparisons of gamma power responses at individual peak gamma frequency (see Table S1) between TACS and Sham trials. Right columns contain t-statistics and p-values for paired-sample t-tests. For all stimulation frequencies, occipital but not frontal control TACS caused a significant reduction of visual stimulus-induced gamma power.

**Paired Samples T-Tests**
|  | t | df | p |
| --- | --- | --- | --- |
| Occipital: IAF - 4 Hz - Sham: IAF - 4 Hz | -6.034 | 14 | < .001 |
| Occipital: IAF - Sham: IAF | -4.317 | 14 | < .001 |
| Occipital: IAF + 4 Hz - Sham: IAF + 4 Hz | -3.112 | 14 | .008 |
| Frontal: IAF - 4 Hz - Sham: IAF - 4 Hz | 0.455 | 14 | 0.656 |
| Frontal: IAF - Sham: IAF | -0.919 | 14 | 0.374 |
| Frontal: IAF + 4 Hz - Sham: IAF + 4 Hz | -0.363 | 14 | 0.722 |

*Note.* Student's t-test.

**Figure S1.**
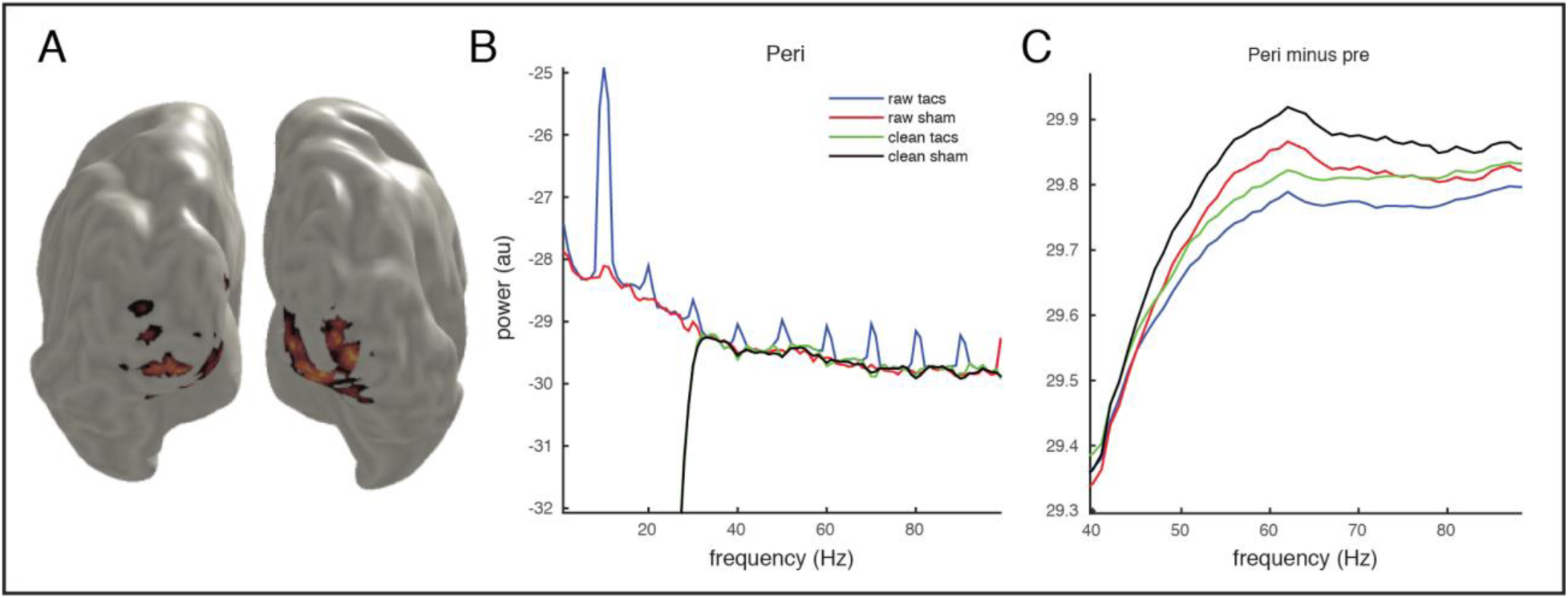
Voxel selection and cleaning. **(A)** Virtual channels were extracted from the 10 voxels, in the visual cortex, showing the highest increase in gamma-band power from baseline in response to visual stimulation. Warm colors in the transversal slice indicate areas showing increased gamma-band activity due to visual stimulation. **(B)** Power spectrum (0.5-1.5 s peri-stimulus period, zero-padded to 10 seconds, Hanning taper) for a representative subject from a single 10 Hz TACS trial before (Raw; blue and red trace) and after (FFT interpolated; green and black trace) artifact suppression. **(C)** Same as in B, but for the difference between peri- and pre-stimulus perods (multitapered, as in TFR). The absence of visible artifacts at the harmonics of the stimulation frequency suggests that the peri-pre subtraction approach and the high-pass filter were already able to remove most of the artifacts (even without FFT interpolation.

**Figure S2.**
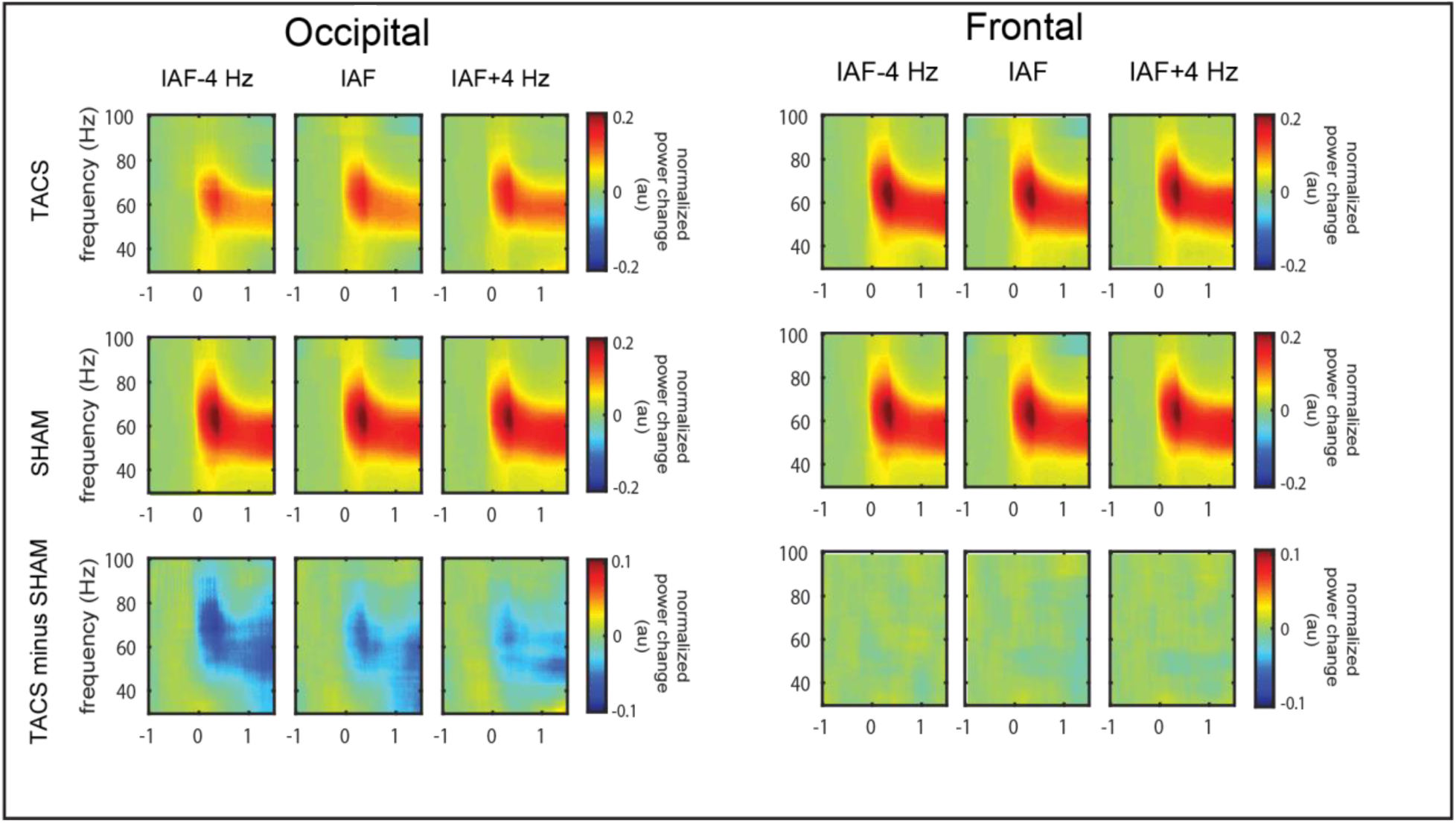
Occipital TACS suppresses visually-induced gamma-band response. TFRs show the visually-induced gamma-band responses for occipital (left panel) and frontal (right panel) TACS, as well as for all stimulation frequencies (columns), including corresponding Sham trials. The bottom row represents the difference between respective TACS and Sham trials.

**Figure S3.**
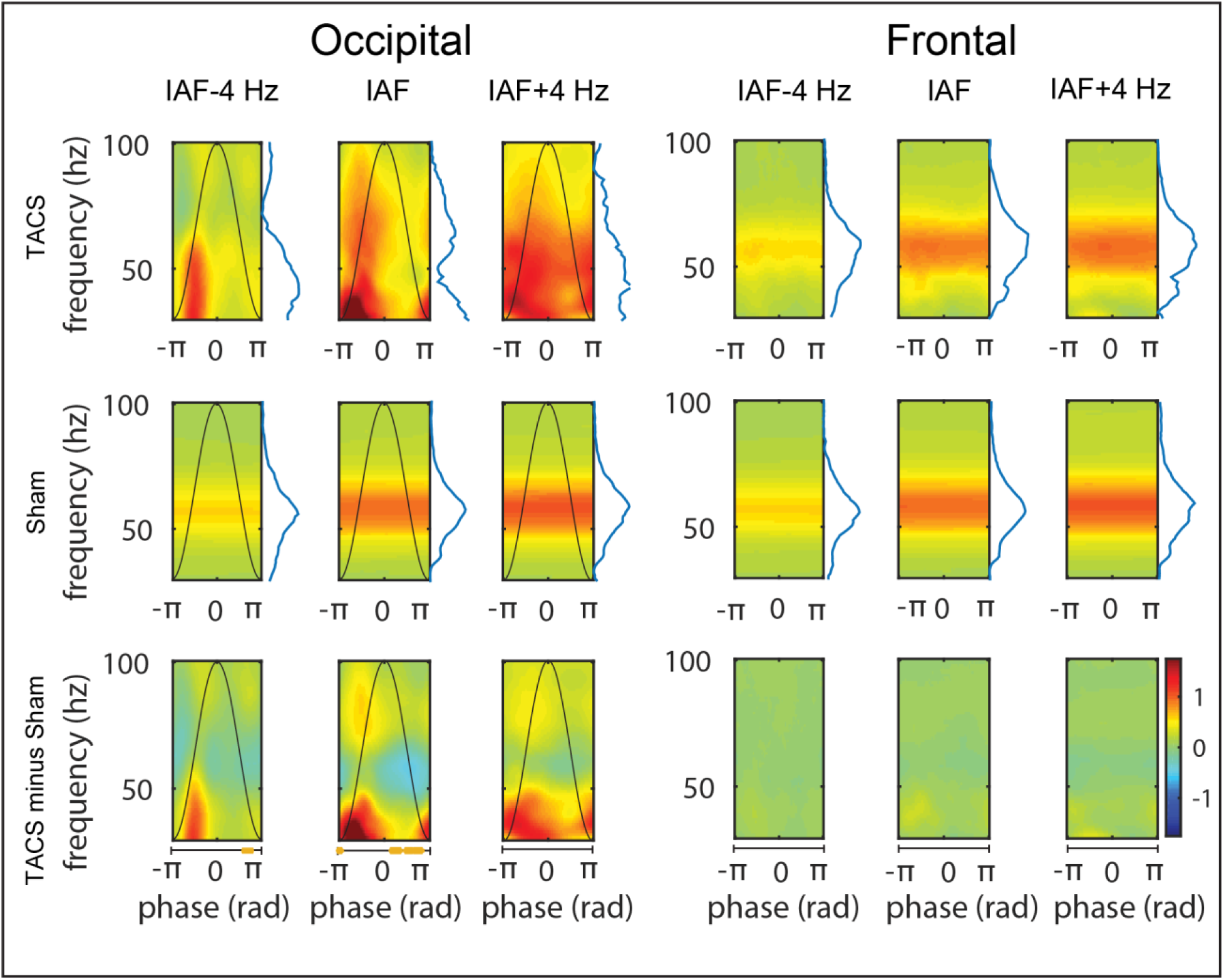
Occipital TACS phasically modulates visually-induced gamma-band response. TACS peak-locked TFRs show gamma-band amplitude sorted according to phase of TACS. Note that some of the apparent differences between TFRs for different frequencies (cf. columns of middle row, representing Sham trials) are partially due to the different cycle length (i.e. time window) that had been normalized into radians as longer segments are accompanied with less frequency smoothing. However, respective Sham trials have been treated the same way and comparisons to Sham thus correct for this effect. Only occipital stimulation shows a clear modulation of gamma-band amplitude according to TACS phase (orange bars underneath TFR indicate significant difference from zero, p_FDR_ < 0.05). The bottom row represents the difference between the TACS and Sham trials.

**Figure S4.**
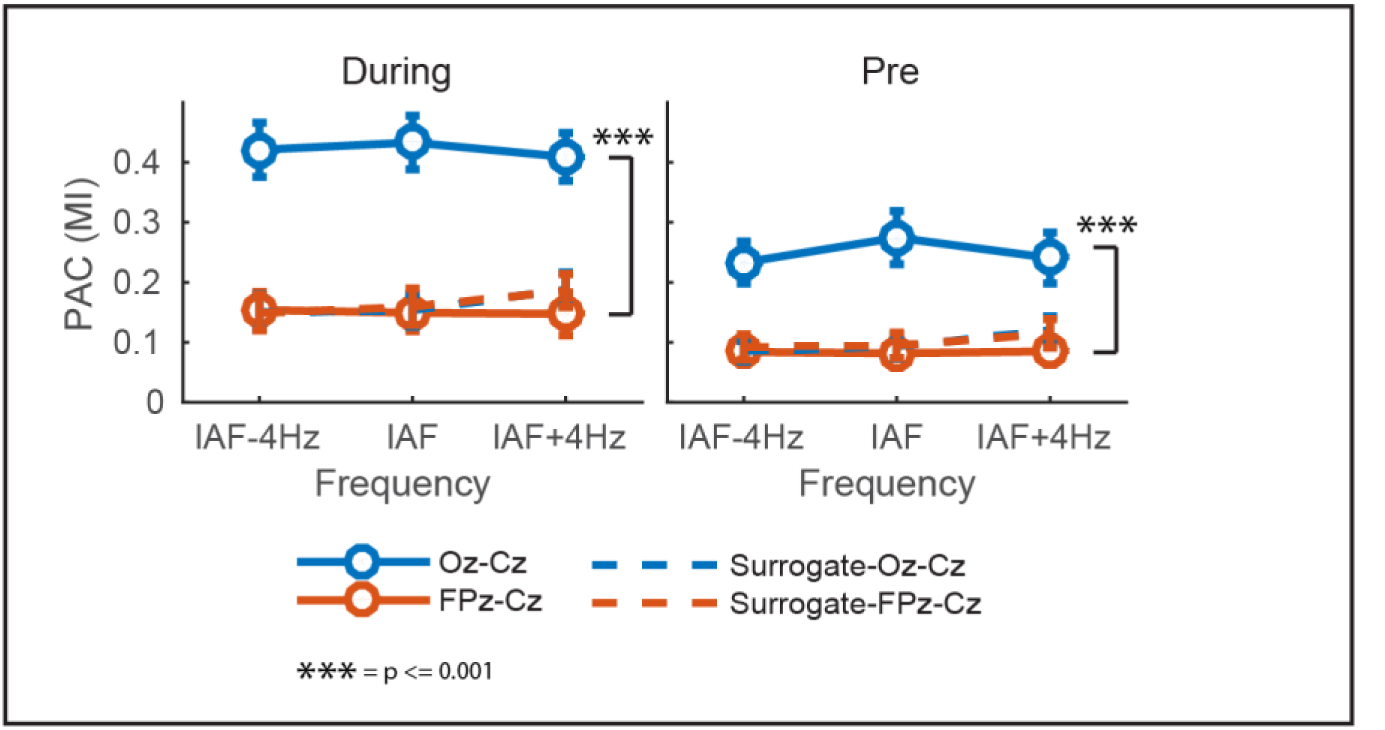
Gamma-band activity phase-coupled to TACS for occipital stimulation. Both during visual stimulation (left) and to a smaller degree also before visual stimulus onset (right), modulation index (MI) analyses revealed increased phase-amplitude coupling (PAC) for occipital TACS relative to both frontal control TACS and surrogate data, whereas PAC for frontal TACS did not differ from surrogate PAC.

